# Branched-chain α-keto acid metabolism is controlled by SLC25A35-mediated mitochondrial import

**DOI:** 10.64898/2026.11.14.766690

**Authors:** Yann Deleye, Jacquelyn M. Walejko, Iuliaa Karavaeva, Neha Majety, Sreya Varanasi, Sean Ferrano, Esra Akkaya, Wentao He, Matthew W. Foster, Olga Ilkayeva, Guo-Fang Zhang, Zachary Gerhart-Hines, Robert W. McGarrah, Phillip J. White

**Affiliations:** Duke Molecular Physiology Institute, Duke University School of Medicine; Durham, NC, USA; Novo Nordisk Foundation Center for Basic Metabolic Research, University of Copenhagen; Copenhagen, DK; Duke Proteomics and Metabolomics Core Facility, Duke University School of Medicine; Durham, NC, USA; Department of Medicine, Division of Endocrinology, Metabolism, and Nutrition, Duke University School of Medicine; Durham, NC, USA; Center for Adipocyte Signaling (ADIPOSIGN), University of Southern Denmark; Odense, DK; Department of Medicine, Division of Cardiology, Duke University School of Medicine; Durham, NC, USA

## Abstract

Liver is considered a primary site for systemic branched-chain α-keto acid (BCKA) oxidation due to its abundant dehydrogenase content. However, hepatocytes rely entirely on import of substrate generated elsewhere because they lack intrinsic branched-chain transaminase activity. The mechanisms governing BCKA transit into the mitochondrial matrix remain unknown. We found that mitochondrial import, rather than dehydrogenase activity, determines hepatocellular capacity for BCKA oxidation. We identified MCT1 and MCT2 as transporters required for cellular entry and SLC25A35 as the essential mitochondrial BCKA carrier. Crucially, hepatic SLC25A35 expression is low and limits the liver’s contribution to systemic clearance. These findings establish the molecular machinery for BCKA transport and redefine the liver’s role in BCKA homeostasis, revealing that mitochondrial transporter abundance, rather than dehydrogenase levels, dictates BCKA metabolic flux.

## INTRODUCTION

The branched-chain amino acids (BCAAs)—leucine, isoleucine, and valine—comprise approximately 45% of the essential amino acid pool (*1*). BCAA metabolism heavily influences systemic nitrogen balance and integrates directly with glucose, lipid, and protein turnover (*1–3*). A central feature of BCAA metabolism is the interconversion and trafficking of BCAA-derived nitrogen and α-ketoacid carbon intermediates across organelle, cell, and tissue compartments. During fasting, for example, mitochondrial branched-chain aminotransferase (BCAT2) deaminates BCAAs in skeletal muscle; where the resulting glutamate serves as the primary nitrogen donor for pyruvate-to-alanine conversion in the glucose-alanine cycle (*4–6*). Concurrently, the branched-chain α-keto acid (BCKA) byproducts are dissipated from the mitochondrial matrix through a combination of export to the systemic circulation and local oxidation via the BCKA dehydrogenase (BCKDH) enzyme complex (*7*). Because BCAT2 activity far outweighs BCKDH capacity in skeletal muscle (*1,8*), BCKAs are preferentially exported to distal tissues where they may undergo further catabolism or be recycled to BCAA (*7,9*). The irreversible decarboxylation of BCKA by BCKDH is considered the first rate-limiting step in BCAA catabolism, generating an array of cell- and context-specific metabolites with distinct biological functions (*1–3*). These products include the TCA cycle intermediates acetyl-CoA and succinyl-CoA, the ketone acetoacetate, the myokine L-β-aminoisobutyric acid (L-BAIBA), the paracrine factor 3-hydroxyisobutyrate (3-HIB), odd-chain monomethyl branched-chain fatty acids, and heme (*7–13*). The liver exhibits among the highest intrinsic BCKDH activities and is thought to serve as a major site for BCKA oxidation. Yet, in stark contrast to skeletal muscle, hepatocytes express virtually no BCAT2. This creates an isolated enzyme system entirely reliant on importing circulating BCKAs into the mitochondrial matrix (*1,14*). Surprisingly, the specific transport mechanisms governing BCKA channeling across cellular and mitochondrial membranes have not been established.

## RESULTS

### Activating BCKDH in liver does not alter BCKA levels in liver or blood of mice

The activity of the BCKDH complex is controlled by enzyme abundance and the level of inhibitory phosphorylation on serine 293 of the E1a subunit. Phosphorylation on serine 293 is determined by the balance between the BCKDH kinase, BDK, and the phosphatase, PPM1K (*14–15*). To quantify the impact that modulation of BCKDH activity in liver exerts on systemic BCKA homeostasis we generated mice in which BDK, the kinase that phosphorylates and inhibits BCKDH, is selectively deleted in hepatocytes (BDK^LKO^). We then compared the effects of hepatic BDK deletion to wild-type (WT) littermates treated with vehicle or the small molecule inhibitor of BDK, 3,6-dichlorobenzo[*b*]thiophene-2-carboxylic acid, BT-2, which lowers BCKDH phosphorylation systemically (*16*). BT-2 administration lowered BCKDH phosphorylation in liver by approximately 50% compared to vehicle treated WT mice (Fig. 1A) and led to a significant reduction in all three BCKA in plasma but did not alter their abundance in liver (Fig. 1B-C). Surprisingly, although hepatic BDK deletion completely abolished the inhibitory phosphorylation present on BCKDH in liver (Fig. 1A), the concentration of BCKA in plasma and liver of BDK^LKO^ mice was not different from vehicle treated WT littermates (Fig.1B-C). There was also no effect of liver BDK deletion on plasma BCAA levels (data not shown). These data reveal that activating the BCKDH complex in liver has no impact on systemic BCKA homeostasis in healthy mice and indicate that other sites of action must be responsible for the BCKA lowering effect of the small molecule inhibitor of BDK, BT-2.

**Fig. 1.**
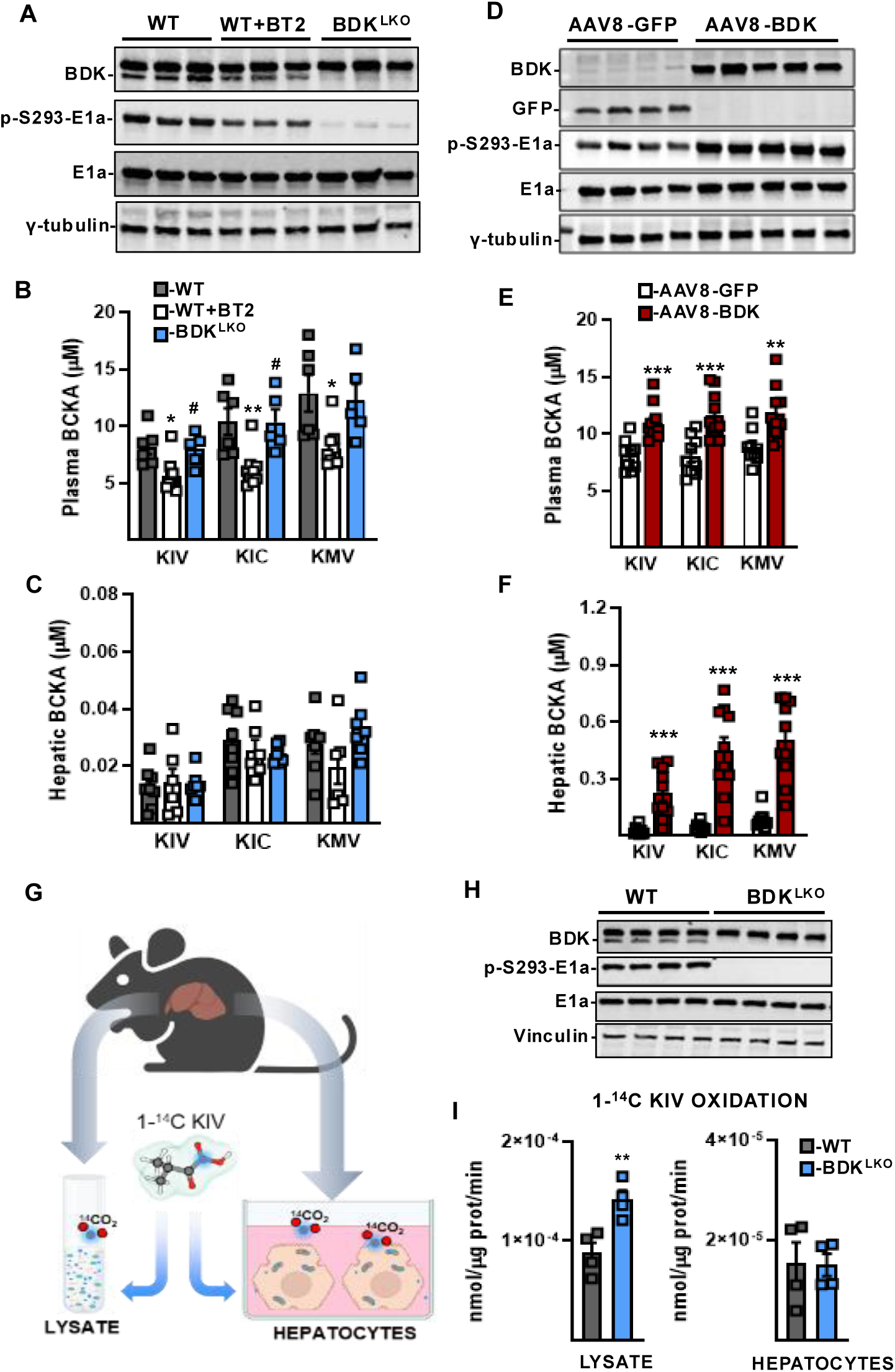
Membrane transport limits hepatic BCKA decarboxylation capacity. (**A-C**) BDK fl/fl mice were treated with either 20mg.kg^-1^ BT2 or DMSO vehicle for seven consecutive days and compared to their liver specific BDK knockout littermates. (A) Western Blot analysis of BDK, phosphorylated Ser293 of Bckdh-E1a (E1a), E1a total and γ-tubulin in the liver. (B) Plasma and (C) hepatic concentrations of branched-chain α-keto acids (BCKA). (**D-F**) C57Bl6 male mice received a single injection of AAV-8-CMV-GFP (n=9) or AAV8-CMV-BDK (n=10). (D) Western Blot analysis of BDK, GFP, phosphorylated E1a, total E1a and γ-tubulin protein levels in the liver. (E) Circulating and (F) hepatic concentrations of BCKA. (**G-I**) Mouse primary hepatocytes were isolated from BDK LKO mice and their littermate controls (G) Schematic of 1- [^14^C]-KIV oxidation assay in lysed or live primary hepatocytes. (H) Western Blot analysis of BDK, phosphorylated and total E1a in primary hepatocytes. (I) Left panel, rate of 1-[^14^C]-KIV oxidation in the lysate and right panel in live primary hepatocytes from WT and BDK ^LKO^ mice, n=4 per group. Statistics: All values are expressed as means ± SEM. (B) One-way ANOVA with Tukey’s multiple comparisons test, * compared to WT, # compared to WT+BT2, * or # p <0.05, ** p <0.01. (E,F,I) Unpaired two-sided t-test, ** p <0.01, *** p <0.001.

### Hepatic BCKDH inhibition augments BCKA in liver and blood without altering BCAA

Since hepatic BDK deletion eliminated BCKDH phosphorylation in liver without impacting BCKA levels (Fig 1A-C), we next tested whether raising BDK expression above baseline in liver has any effect on BCKA homeostasis. We overexpressed recombinant BDK in liver of C57Bl6 mice using adenoassociated-virus serotype-8 (AAV8) in which human BDK was expressed on a cytomegalovirus (CMV) promoter. An AAV8-CMV construct expressing green-fluorescent protein (GFP) was used as the control. Retro-orbital administration of AAV8-BDK led to a robust induction of BDK in liver as well as heightened inhibitory phosphorylation on BCKDH (Fig. 1D) but did not increase BDK levels or change the phosphorylation state of BCKDH in muscle, heart, or perigonadal white adipose tissue (pgWAT) compared to AAV8-GFP treated control mice (fig. S1A-C). This effect on liver BCKDH phosphorylation was accompanied by a 30% increase in plasma BCKA and 3-fold higher BCKA levels in liver 8-weeks post AAV8-BDK administration compared to AAV8-GFP treated mice (Fig. 1E-F). Consistent with the extremely low levels of BCAT2 in liver, the effect of AAV8-mediated hepatic BDK overexpression to raise plasma and liver BCKA occurred without any effect on circulating BCAA (fig. S1D).

### Substrate supply determines the upper threshold for BCKA oxidation in liver

We reasoned that the seemingly opposing findings from our liver BDK knockout and overexpression studies could be explained by a model wherein substrate supply rather than BCKDH activity per se determined the upper threshold for BCKA oxidation in liver. In this model, inhibitory BCKDH phosphorylation exerts control over BCKA oxidation only when it causes enzyme activity to fall below the limit of substrate provision.

To directly evaluate the influence of membrane transport on BCKDH-mediated BCKA oxidation in hepatocytes, we quantified the impact of BDK deletion on BCKA oxidation in primary hepatocytes in which membranes were intact and compared the outcome to those obtained in parallel assays using liver tissue lysates in which the membranes no longer provide a functional barrier for substrate access to BCKDH (Fig. 1G). Liver lysates and primary hepatocytes from WT or BDK^LKO^ mice were incubated with the 1-[^14^C]-labeled ketoacid derivative of valine, α-ketoisovalerate (KIV), and the^14^CO_2_ generated by BCKDH mediated decarboxylation of 1-[^14^C]- KIV was subsequently trapped and counted (*15*). Whereas liver lysate from BDK^LKO^ mice displayed higher BCKDH activity compared to WT mice (Fig. 1I), primary hepatocytes cultured from the same mice, displayed rates of BCKA oxidation that were not different to their WT counterparts (Fig. 1I). Notably, as observed in liver lysates (Fig. 1A), the inhibitory phosphorylation on ser293 of BCKDH was undetectable in hepatocytes derived from BDK^LKO^ and robust in hepatocytes from WT mice (Fig. 1H). Similar results were also obtained with BT2-treated liver lysates and hepatocytes from C57Bl6 mice (fig. S1E-F). Together these experiments support the substrate provision model of hepatocyte BCKA oxidation wherein membrane transport of BCKA rather than BCKDH active state determines the upper threshold for BCKA oxidation in liver of BDK^LKO^ and BT2-treated mice.

### Monocarboxylic acid transporter activity is required for systemic BCKA homeostasis

The mechanisms governing systemic BCKA trafficking across tissues have not been defined, however, the monocarboxylate carriers have been implicated in the neuron to astrocyte cycling of BCKA that supports cytosolic glutamate synthesis for its neurotransmitter function (*17*). We therefore assessed whether AR-C155858, a small molecule inhibitor of MCT1 and MCT2, alters systemic BCKA homeostasis and BCKA clearance in C57Bl6 mice. Mice were fasted for 4 hours then administered vehicle or AR-C155858 at a dose of 5mg/kg intraperitoneally. Ninety minutes following the administration of the MCT1/2 inhibitor a baseline plasma sample was taken, after which a 100mg/kg bolus of the three BCKA was administered into the intraperitoneal cavity of the mice. BCKA were then assayed in plasma samples taken at 5, 15, and 45 minutes (Fig. 2A). Here AR-C155858 administration resulted in a significant increase in the known MCT substrate β-hydroxybutyrate (β-HB; fig. S2A) as well as the levels of all three BCKA in blood (Fig. 2B). Consistent with a specific effect on BCKA transport, levels of the three cognate BCAA were unaffected by AR-C155858 administration (fig. S2B). Clearance of the α-keto acids derived from valine and leucine, α-ketoisovalerate (KIV) and α-ketoisocaproate (KIC), was also less efficient following the BCKA bolus in mice that received AR-C155858 (Fig. 2C-D). However, AR-C155858 administration had no impact on clearance of the α-keto acid from isoleucine, α-ketomethylvalerate (KMV), which displayed substantially slower kinetics than KIV and KIC in control mice (Fig. 2E).

**Fig. 2.**
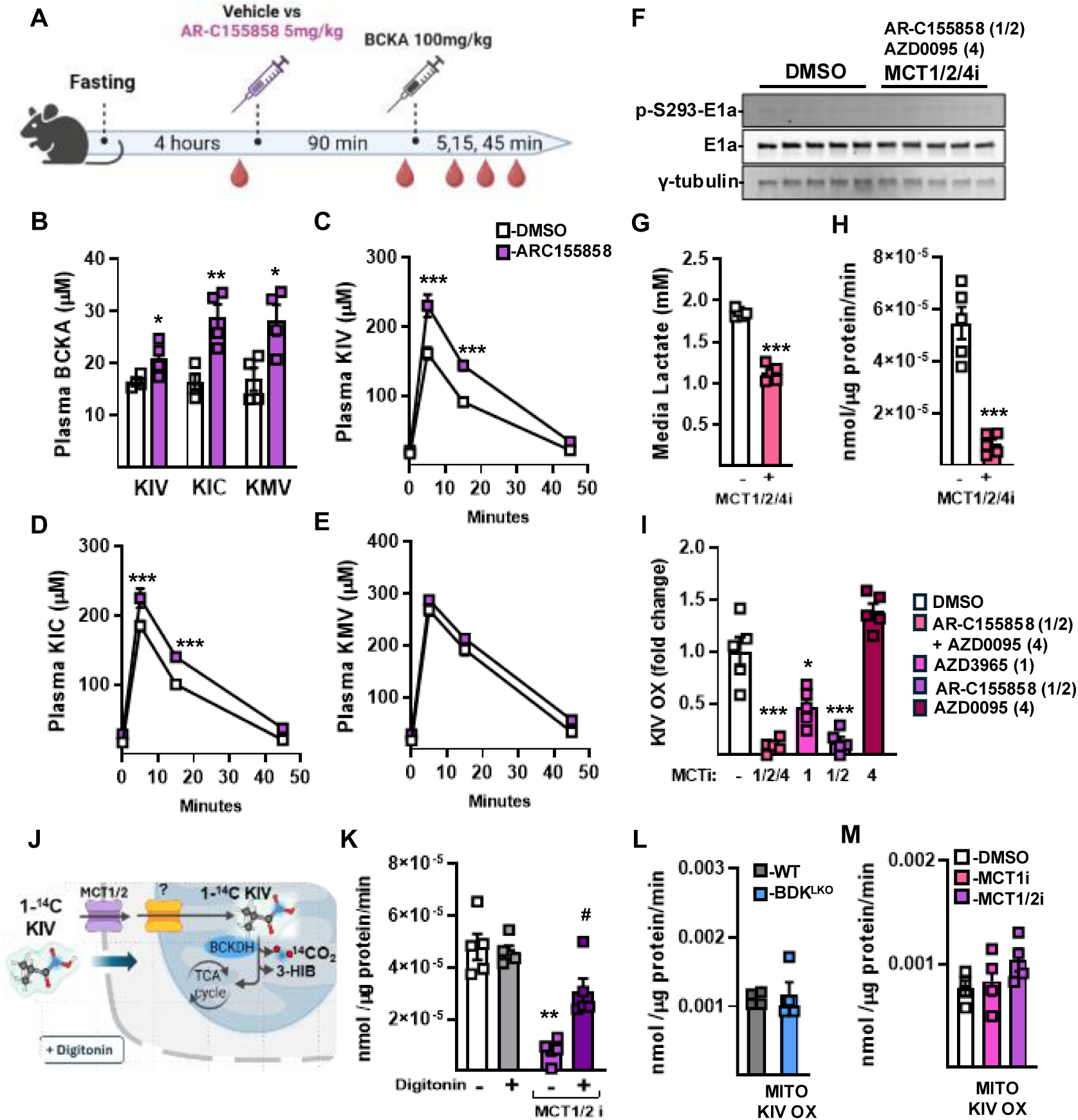
MCT1/2 transport shapes systemic BCKA homeostasis and is required for BCKA metabolism in intact hepatocytes but not mitochondria. (A-E) C57Bl6 male mice were fasted for 4h and received a single intraperitoneal injection of AR-C155858 (MCT1/2 inhibitor; 5mg/kg) or DMSO, 90 minutes prior to a bolus of 100mg.kg^-1^ BCKA (n=4 animals per group). (A) Study design. (B) Circulating BCKA 90min post MCT1/2 inhibitor injection. Circulating (C) KIV, (D) KIC, (E) KMV concentrations during the BCKA tolerance test. (**G-I**) Primary hepatocytes were isolated from BDK^LKO^ male mice and incubated for one hour with different combination of MCT1,2 and/or MCT4 inhibitors. (G) Lactate concentration in the media from primary hepatocytes incubated with MCT1,2,4 inhibitors. (H) Rate of 1-[^14^C]-KIV oxidation in primary hepatocytes treated with a cocktail of MCT1/2 and 4 inhibitors (AR-C155858 250nM + AZD0095 250nM) or (I) various inhibitors for either MCT1 (AZD3965; 100nM), MCT1/2 (AR-C155858, 250nM), MCT4 (AZD0095; 250nM) or MCT1/2, and 4. (**J**) Schematic of 1-[^14^C]-KIV oxidation in live permeabilized primary hepatocytes treated with digitonin (7.5µg/mL) and (**K**) rate of 1-[^14^C]-KIV oxidation in intact or membrane permeabilized primary hepatocytes treated with MCT1/2 inhibitor (AR-C155858, 250nM) or DMSO. (**L**) Rate of 1-[^14^C]-KIV oxidation in isolated hepatic mitochondria from WT and BDK^LKO^ mice. (**M**) Rate of 1-[^14^C]-KIV oxidation in isolated hepatic mitochondria from WT mice treated with MCT1 inhibitor (AZD3965; 100nM), MCT1/2 inhibitor (AR-C155858; 250nM) or DMSO. *Statistics:* All values are expressed as mean ± SEM. (B, G, H) Unpaired two-tailed t-test, * p <0.05, ** p <0.01, *** p <0.001. (C, D) Two-way ANOVA with Šidák multiple comparisons test, *** p <0.001. (I) One-way ANOVA with Dunnett’s multiple comparisons test, *** p <0.001. (K) One-way ANOVA with Tukey’s multiple comparisons test, # p <0.05 compared to MCT1/2i without digitonin, ** p < 0.01 compared to DMSO.

### MCT1 and MCT2 are required but not rate limiting for BCKA oxidation in hepatocytes

We next directly assessed the impact of MCT inhibition on BCKA oxidation in primary hepatocytes. Here we incubated primary hepatocytes from BDK^LKO^ mice with a cocktail containing the MCT1/2 inhibitor, AR-C155858, and the MCT4 inhibitor, AZD0095. Hepatocytes from BDK^LKO^ mice were employed for this initial study as they have no inhibitory phosphorylation on BCKDH and would therefore be refractory to any potential off target effect to increase the levels of BDK-mediated BCKDH phosphorylation which could confound the results. Incubation of primary hepatocytes with the MCT1/2/4 inhibitor cocktail did not alter BCKDH phosphorylation or abundance (Fig. 2F) but did lower media lactate indicative of the expected inhibition of hepatocyte MCT activity (Fig. 2G). Remarkably, the MCT1/2/4 inhibitor cocktail also completely abolished BCKA oxidation in primary hepatocytes (Fig. 2H). Thus, MCT activity is required for the oxidation of extracellular BCKA by hepatocytes.

To determine which MCT isoforms contribute to BCKA transport we repeated the study in primary hepatocytes from C57Bl6 mice and compared the effect of the MCT1/2/4 inhibitor cocktail with AZD3965 which inhibits MCT1 selectively, AR-C155858 which inhibits only MCT1 and MCT2, and AZD0095, which inhibits MCT4 selectively (Fig. 2I). As AZD3965 and AR-C155858 significantly impaired BCKA oxidation but AZD0095 did not, these experiments provided support for a role for both MCT1 and MCT2 in BCKA transport into hepatocytes and suggested that MCT4 does not contribute to BCKA transport in this cell type (Fig. 2I).

The above studies revealed that exposure of hepatocytes and mice to MCT1 and MCT2 inhibitors impairs BCKA metabolism; however, they did not provide evidence that this effect is directly related to membrane transport or clarify whether plasma membrane transport is rate limiting for BCKA oxidation in hepatocytes. We therefore performed a BCKA oxidation study in which the plasma membrane of the primary hepatocytes was left intact or permeabilized with digitonin (Fig. 2J). In this experiment, digitonin did not alter the level of BCKA oxidation in vehicle treated cells but restored BCKA oxidation to near control levels in hepatocytes treated with the MCT1/2 inhibitor (Fig. 2K). Importantly, neither digitonin nor the MCT1/2 inhibitor altered BCKDH phosphorylation or abundance (fig. S2C). Thus, these data indicate that activity of MCT1 and MCT2 is required for BCKA transport across the cell membrane. However, since permeabilizing the membrane with digitonin did not raise oxidation compared to vehicle it does not appear that plasma membrane transport determines the upper threshold for BCKA oxidation in hepatocytes.

### Mitochondrial BCKA import restrains BCKA oxidation in liver

Given the results of the digitonin experiment, we next purified liver mitochondria from WT and BDK^LKO^ mice and repeated the BCKA oxidation assay to determine whether the mitochondrial membrane restrains BCKA oxidation in hepatocytes. As observed in primary hepatocytes, loss of BDK completely abolished inhibitory phosphorylation on BCKDH (fig. S2D) yet had no effect to increase BCKA oxidation in isolated mitochondria (Fig. 2L). Similar results were obtained in mitochondria incubated with BT-2 (fig. S2E-F). However, consistent with the findings from our study of AAV8-mediated BDK overexpression, mitochondria from livers of mice lacking the BCKDH phosphatase, PPM1K (PPM1K^LKO^), displayed both higher BCKDH phosphorylation and lower levels of BCKA oxidation. These data indicate that BCKA trafficking from the cytosol into the mitochondrial matrix is likely rate-limiting for BCKA oxidation in hepatocytes.

To rule out the possibility that the MCT1/2 inhibitor cocktail was acting on the carriers in the mitochondria in addition to the cell membrane we also conducted a BCKA oxidation experiment in liver mitochondria from C57Bl6 mice incubated with the MCT1 inhibitor or the MCT1/2 inhibitor. Importantly, as opposed to their robust inhibitory effect on BCKA oxidation in primary hepatocytes neither MCT inhibitor was found to have any impact on BCKA oxidation in isolated mitochondria (Fig. 2M). These data indicate that BCKA trafficking from the cytosol to the mitochondrial matrix does not involve the actions of MCT1 or MCT2.

### Bioinformatics approach identifies a candidate mitochondrial BCKA carrier

The results obtained thus far suggest that BCKA transport into the mitochondrial matrix rather than across the cell membrane dictates the upper threshold for BCKA oxidation in hepatocytes. Transport of mitochondria-compartmentalized metabolites relies on membrane-bound carrier proteins in the inner mitochondrial membrane (IMM). The SLC25 solute carrier family encodes the mitochondrial solute carriers present in the IMM and has 53 members. Whereas carriers have been defined for multiple mitochondrial metabolites there is no known carrier for BCKA in the IMM and there remains multiple orphan carriers with no defined substrates (*18*). Given that mitochondrial transport is required for BCKDH activity, we reasoned that the mitochondrial solute carrier for BCKA likely co-evolved alongside BCKDH. We therefore applied a phylogenetics analysis using the PhyloGene server to determine whether any mitochondrial solute carriers co-evolved alongside the alpha subunit of BCKDH (*19*). This analysis highlighted five members of the SLC25 family among the top twenty genes found to co-evolve with *Bckdha* (Fig. 3A). Of the five carriers in the list, all except SLC25A35 had defined substrates (*20*). SLC25A27 transports protons across the IMM and is commonly referred to as uncoupling protein-4; SLC25A2 and SLC25A15 are mitochondrial ornithine carriers, ORNT2 and ORNT1 respectively; and SLC25A47 is a mitochondrial carrier for the BCKDH cofactor NAD^+^.

**Fig. 3.**
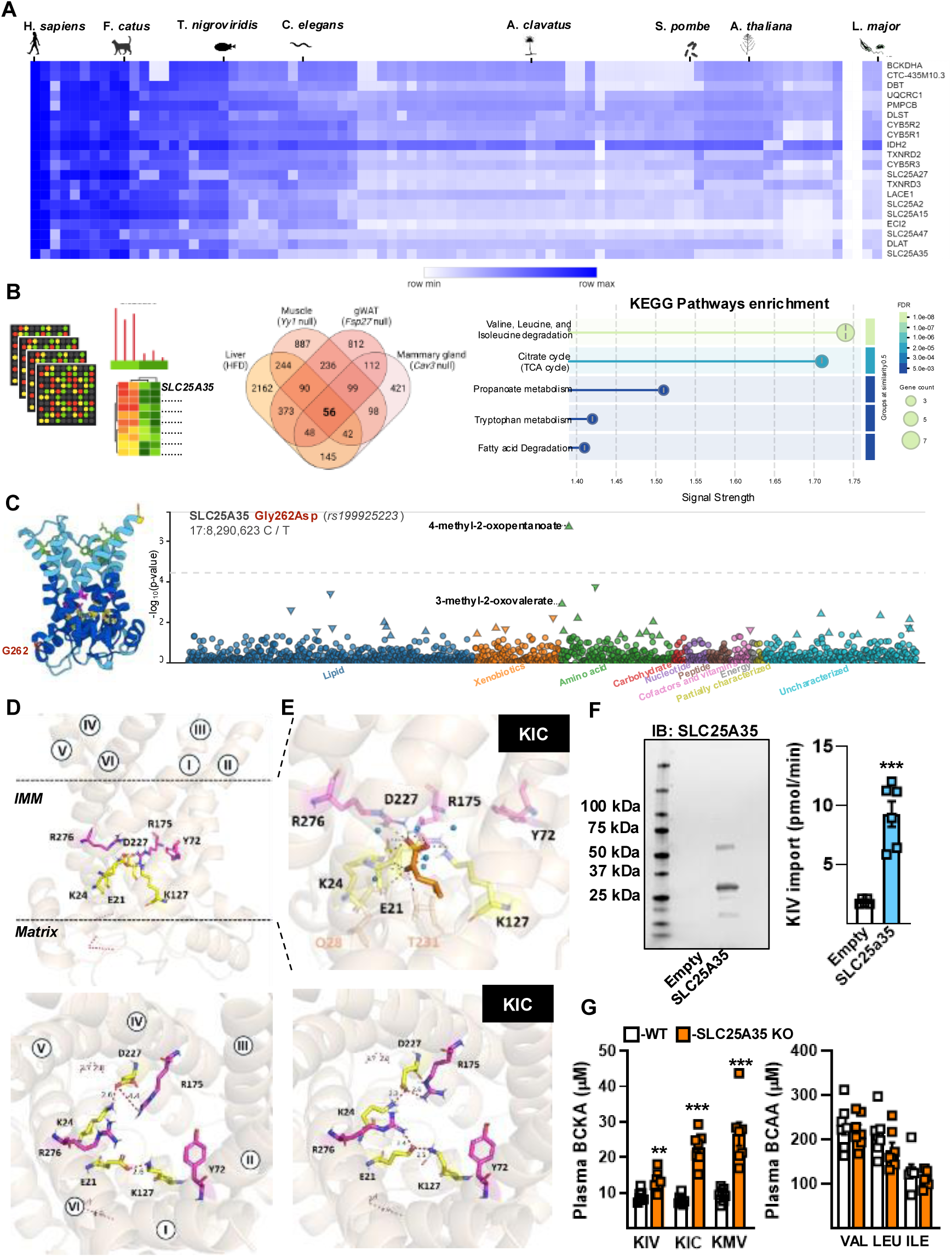
Integrative genomics, structural analysis, and functional validation identify SLC25A35 as a mitochondrial BCKA transporter. (**A**) Heatmap from Phylogene co-evolution analysis showing top 20 genes that co-evolved with, BCKDHA, the alpha subunit of BCKDH. (**B**) From left to right of panel, workflow diagram, Venn diagram, and pathways enrichment results for genes co-regulated with *Slc25a35* in four publicly available datasets in which *Slc25a35* expression was significantly up- or down-regulated. (**C**) Left panel – Cartoon representation of a lateral view of the Alphafold predicted structure of human SLC25A35 in the c-state where the matrix gate is in the closed position. Residues in the common binding sites are colored in pink; residues that form the matrix salt bridge network are colored in yellow, and residues that form the cytoplasmic salt bridge network are shown in green. Glycine G262 is colored in red. Right panel – Manhattan plot of plasma metabolite associations for 1,390 serum metabolites with the SNP rs199925223 in SLC25A35 (Gly262>Asp) in 6,136 Finnish people from the METSIM phenome-wide association study. (**D, E**) Modeling of KIC docking using the Alphafold predicted structure of SLC25A35 in the c-state. Panels in (D) – show SLC25A35 in the unbound state. Panels in (E) – show SLC25A35 upon simulated hydrated-flexible docking of KIC, represented in the upper panel in orange stick form and not shown in the lower panel. The 6 transmembrane α-helices are colored in wheat. Salt bridge networks and interactions (red dashed lines) are shown between residues from the common binding sites (pink sticks) and residues of the matrix gate (yellow sticks). Polar interactions are represented with dashed grey lines. Other non-flexible residues found to form interactions with KIC are shown in wheat. In both (D) and (E), top panels – show SLC25A35 from the lateral view. Bottom panels – show SLC25A35 from the cytosolic side of the inner mitochondrial membrane (IMM) with distances between residues of the common binding sites and matrix salt bridge network indicated in Angstroms. (**F**) SLC25A35 western blot (left panel) and KIV import rate (right panel) for liposomes containing recombinant SLC25A35 and control empty liposomes. (**G**) Plasma BCKA and BCAA concentrations from SLC25A35 knock-out mice and their wild-type littermates. *Statistics:* All values are expressed as mean ± SEM. (F, G) Unpaired two-tailed t-test, **P<0.01, *** P<0.001.

As SLC25A35 was the only carrier without a defined substrate on the list, we next performed a co-expression analysis to determine which biological pathways are dynamically regulated in coordination with SLC25A35. For this analysis, we identified four independent gene sets in the GEO gene expression database in which SLC25A35 was significantly up or downregulated and filtered for genes that were significantly regulated in the same direction as SLC25A35 (Fig. 3B). This analysis yielded a set of 56 genes that were co-regulated with *Slc25a35* expression across all four data sets (Fig. 3B). Remarkably pathway analysis of the *Slc25a35* co-regulated gene-set identified the valine, leucine, isoleucine degradation pathway as the most enriched pathway (Fig. 3B), with seven genes *Dbt*, *Ivd*, *Acadsb*, *Hibadh*, *Aldh6a1*, *Aox1*, and *Pcca* displaying co-regulation with *Slc25a35* (fig. S3A). Together these co-evolution and co-expression analyses pointed to the orphan solute carrier SLC25A35 as a strong candidate for mitochondrial BCKA transport.

### Human genetics links SLC25A35 to BCKA levels

To substantiate the association between SLC25A35 and BCKA, we next explored metabolite associations with the *SLC25A35* locus using data from 6,136 Finnish persons in the PheWeb METSIM metabolomics portal (*21*). Remarkably, of the 1,391 plasma metabolites that were quantified, the strongest metabolite signal for the *SLC25A35* region was elevated levels of KIC, the α-ketoacid from leucine, annotated as 4-methyl-2-oxopentanoate in METSIM (Fig. 3C). This relationship was driven by a single coding variant in *SLC25A35* (rs199925223) that was strongly associated with higher plasma KIC (P=1.7 x 10-7) and nominally associated with elevated plasma KMV (P=1.1 x 10-3). The rs199925223 variant is a cytosine to thiamine substitution that changes codon 262 of *SLC25A35* from glycine to aspartate. This glycine residue forms a flexible hinge connecting the transmembrane helix H6 with the matrix helix H56 and is highly conserved among members of the SLC25 family (fig. S3B). Notably, site-directed mutagenesis of the glycine residue at position 262 to aspartate has been shown to completely abolish SLC25 carrier activity *in vitro* (*22*). Together these data provide human genetic support for SLC25A35 as a mitochondrial carrier for BCKA import.

### *In silico* modeling supports SLC25A35-mediated transport of BCKAs

To determine if BCKAs could serve as substrates for SLC25A35, we conducted docking simulations of the three individual BCKAs, KIC, KIV, and KMV, with an AlphaFold-predicted human SLC25A35 structure using AutoDock Vina (*23, 24*). Simulations incorporating a hydrated substrate, flexible binding pocket residues, and a flexible matrix salt-bridge network (Fig. 3D) demonstrated that all three BCKAs dock within the common substrate-binding region of SLC25A35 (Fig. 3E and figs. S3, C and D). Binding affinities were robust and comparable across all three substrates, yielding docking scores of −12.25 kcal/mol for KIC, −12.26 kcal/mol for KIV, and −12.32 kcal/mol for KMV. In the preferred binding poses, the carboxyl and/or keto groups of each BCKA formed critical polar contacts with two conserved arginine residues (R175 and R276) within the central binding domain. These structural arrangements permitted the substrate’s keto and/or carboxyl groups to establish disruptive polar interactions with the four core residues of the matrix salt-bridge network (E21, K24, K127, and D227) (Fig. 3E and figs. S3, C and D). Furthermore, ligand docking induced proximity between the salt-bridge network and the substrate-binding domain, favoring inter-residue interactions required to transition the carrier from a matrix-closed to a matrix-open conformation (*18*). Collectively, these structural data indicate that BCKAs dock with SLC25A35 in a manner structurally congruent with the alternating-access mechanism established for the SLC25 mitochondrial transporter family.

### SLC25A35 directly transports BCKAs and is required for systemic clearance

To validate the transport capacity of SLC25A35 for BCKA, we reconstituted purified recombinant human SLC25A35 into liposomes. Because structural modeling indicated that SLC25A35 possesses a weak cytosolic salt-bridge network (fig. S3E), we evaluated substrate import in uniport mode using 0.1µCi of 1-[^14^C]-KIV as a tracer in a transport buffer containing 2.5mM of unlabeled KIV. SLC25A35-containing proteoliposomes exhibited substantially higher rates of KIV uptake than protein-free control liposomes (Fig. 3F), providing direct biochemical evidence that SLC25A35 acts as a BCKA transporter. To determine the *in vivo* necessity of this transport activity, we evaluated plasma metabolic profiles of global SLC25A35 knockout mice (SLC25A35 KO). Whole-body deletion of SLC25A35 dramatically elevated plasma BCKA concentrations without altering systemic BCAA levels (Fig. 3G). Specifically, plasma levels of KIC and KMV increased threefold relative to wild-type controls, whereas KIV levels displayed a significant yet more modest 30% elevation compared to their wild-type counterparts (Fig. 3G). Notably, this pronounced elevation of KIC and KMV relative to KIV mirrored our human genetic association data (Fig. 3C). Together, these findings firmly establish SLC25A35 as a physiologically essential and substrate-selective mitochondrial transporter for systemic BCKA clearance.

### Hepatic SLC25A35 protein abundance limits mitochondrial BCKA oxidation

To assess the relative abundance of SLC25A35, we performed mass spectrometry-based proteomics on purified mitochondria isolated from primary C57Bl6 mouse hepatocytes. Among the 32 SLC25 family members quantified, SLC25A35 was one of the least abundant transporters, clustering within a low-expression group of 13 members (fig. S4A). Notably, SLC25A35 protein expression matched that of the regulatory BCKDH phosphatase PPM1K, which is less than one-tenth as abundant as core BCKDH complex components and downstream catabolic enzymes (Fig. 4A). To evaluate whether this stoichiometric ratio is lower than expected for an oxidative enzyme-associated carrier, we calculated the transporter-to-enzyme ratios for four highly active hepatic mitochondrial pathways: MDH2, OTC, ACADL, and PDH. The SLC25A35-to-BCKDHA ratio was less than one-third of these canonical carrier-enzyme pairs (Fig. 4B). This exceptionally low expression relative to downstream enzymatic capacity indicates that mitochondrial transport may serve as a rate-limiting step for hepatic BCKA oxidation.

**Fig. 4.**
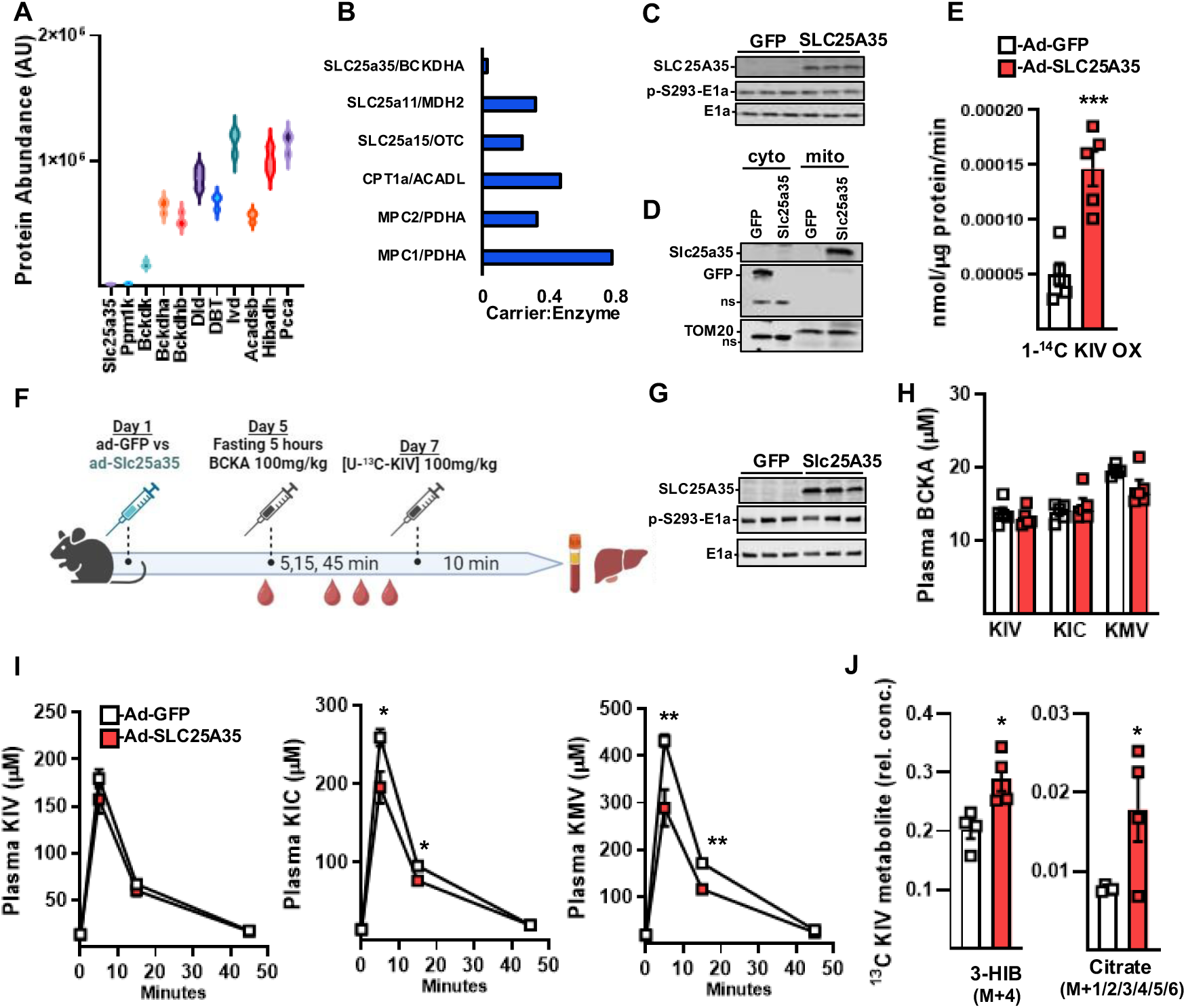
SLC25A35 expression dictates BCKA oxidation and clearance *in vivo*. (**A**) BCAA/BCKA pathway protein abundance in mitochondria isolated from primary hepatocytes, measured by mass spectrometry (n=3) and (**B**) ratio of protein abundance of different carriers with their associated limiting catabolic enzyme. (**C-E**) Primary hepatocytes from C57Bl6 male mice were infected with adenovirus Ad-CMV-SLC25A35 or Ad-CMV-GFP. (C) Western Blot analysis of SLC25A35, Phosphorylated and total E1a protein levels in whole cell lysates, (D) Western Blot analysis of SLC25A35, GFP and TOM20 protein level in cytosol and mitochondrial fractions, and (E) rate of oxidation of 1-[^14^C]-KIV in primary hepatocytes overexpressing SLC25A35 or GFP. (**F-J**) C57Bl6 male mice were injected with Ad-CMV-SLC25A35 (n=5) or Ad-CMV-GFP (n=5). A BCKA tolerance test was performed on day 5 and a bolus of [U-^13^C-KIV] was given 10 min prior euthanasia on day 7. (F) Study Design in mice. (G) Western Blot analysis of SLC25A35, phosphorylated and total E1a in the liver. (H) Circulating BCKA concentration after 5h fasting and circulating (I) KIV, KIC and KMV during a BCKA tolerance test. (J) Mass spectrometry analysis of hepatic concentration of [^13^C]- labeled 3-HIB and citrate, derived from ^13^C-KIV catabolism. *Statistics:* All values are expressed as means ± SEM. (E, J) Unpaired two-tailed t-test, * p <0.05 and *** p <0.001. (I) Multiple unpaired two-tailed t-test, * p <0.05 and ** p <0.01.

To directly test whether baseline SLC25A35 levels restrict BCKA clearance, we transduced primary hepatocytes with adenoviruses driving CMV-promoter-mediated expression of human SLC25A35 (Ad-SLC25A35) or a GFP control (Ad-GFP). Ectopic SLC25A35 expression did not alter BCKDH protein abundance or its inhibitory phosphorylation at Ser293 (Fig. 4C), and the recombinant protein localized exclusively to the mitochondrial fraction (Fig. 4D). Remarkably, Ad-SLC25A35 transduction increased BCKA oxidation rates threefold (Fig. 4E). This metabolic effect was highly specific to SLC25A35 as adenoviral delivery of SLC25A34—the closest sequence homolog to SLC25A35 (*20*)—had no impact on BCKA oxidation (fig. S4, B to D). Collectively, these findings demonstrate that the expression level of SLC25A35, rather than BCKDH enzymatic activity, dictates the upper threshold for hepatic BCKA catabolism.

### Hepatic SLC25A35 overexpression accelerates systemic BCKA clearance and downstream oxidation

Having established that SLC25A35 abundance sets the upper threshold for BCKA catabolism *in vitro*, we next evaluated whether increasing hepatic mitochondrial BCKA transport capacity impacts systemic BCKA homeostasis *in vivo*. We administered Ad-SLC25A35 or Ad-GFP control adenoviruses to C57Bl6 mice, measured five-hour fasting plasma metabolic profiles five days post-injection, and subjected the cohorts to an intraperitoneal BCKA tolerance test (100mg/kg; Fig. 4F). Consistent with our findings in primary hepatocytes, Ad-SLC25A35 treatment robustly increased hepatic SLC25A35 expression without disrupting BCKDH protein abundance or its inhibitory phosphorylation at Ser293 (Fig. 4G). Crucially, no ectopic or off-target SLC25A35 expression was detected in perigonadal white adipose tissue (pgWAT), interscapular brown adipose tissue (BAT), skeletal muscle, or the heart (fig. S4E). Under fasting conditions, hepatic SLC25A35 overexpression did not alter basal plasma levels of BCKAs or BCAAs (Fig. 4H and fig. S4F). However, upon systemic challenge, Ad-SLC25A35-treated mice exhibited significantly enhanced clearance of both KIC and KMV (Fig. 4I). The clearance rate of KIV—the most rapidly cleared endogenous BCKA—was not further accelerated in plasma (Fig. 4I). However, despite the absence of effect on whole animal clearance, *in vivo* stable isotope tracing revealed a profound increase in intrahepatic KIV oxidation following Ad-SLC25A35 treatment. Indeed, mice were administered a bolus of U-[^13^C]-KIV ten minutes prior to euthanasia and liver of Ad-SLC25A35 mice accumulated substantially more KIV-derived mitochondrial metabolites than Ad-GFP controls (Fig. 4J). Notably, KIV-derived [^13^C]-3-HIB concentrations rose by 30%, whereas KIV-derived [^13^C]-citrate levels increased twofold. The more modest accumulation of labeled 3-HIB relative to citrate likely reflects its rapid secretion into circulation as a paracrine factor, whereas citrate is more readily retained within the liver. Collectively, these data demonstrate that hepatic mitochondrial transport is a rate-limiting step for systemic BCKA clearance and subsequent intramitochondrial oxidation *in vivo*.

## DISCUSSION

The discovery of SLC25A35 as an essential mitochondrial BCKA carrier fundamentally redefines the architecture of systemic branched-chain amino acid (BCAA) metabolism. For decades, the irreversible decarboxylation of BCKAs by the mitochondrial BCKDH complex has been universally recognized as the definitive rate-limiting step in BCAA catabolism (*1–3*). Consequently, the liver—possessing the highest relative abundance of active BCKDH complex—was presumed to act as a principal metabolic sink capable of directly clearing systemic BCKAs (*1–3, 8, 14*). However, our findings demonstrate that this classical view overlooks a critical biophysical barrier. Because hepatocytes lack functional BCAT2 activity, they cannot synthesize BCKAs locally (*8*). Instead, they are completely dependent on importing extracellular BCKAs across both cellular and inner mitochondrial membranes. By demonstrating that complete genetic ablation of inhibitory BCKDH phosphorylation in liver fails to lower blood BCKA concentrations, we reveal that downstream dehydrogenase activity operates far below capacity under basal conditions due to limited substrate supply. Our proteomic profiling, *in vitro* transport assays, and *in vivo* BCKA clearance studies reveal that it is the exceptionally sparse mitochondrial expression of the mitochondrial solute carrier SLC25A35 that imposes the true upper threshold on hepatic BCKA oxidation.

By shifting the regulatory paradigm from enzymatic processing capacity to mitochondrial transport kinetics, these insights provide a mechanistic explanation for tracing studies in which BCAA catabolism was not aligned with BCKDH activity (*9*). Our analysis of the METSIM cohort connects a conserved, loss-of-function variant in SLC25A35 (rs199925223) directly to elevated plasma levels of specific BCKAs in humans. Strikingly, this precise biochemical signature is perfectly mirrored in our global SLC25A35 knockout mice, which accumulate circulating KIC and KMV while keeping systemic BCAA levels unperturbed. This substrate selectivity highlights a distinct evolutionary hierarchy among individual BCKAs. The relative protection of KIV clearance from SLC25A35 deficiency could be explained by a more robust transport across the cell membrane. Indeed, whereas loss of SLC25A35 only exerted a modest impact on KIV clearance, this BCKA was the most dramatically altered by MCT inhibition. Together these data indicate that structural differences in alkyl side chains differently impact transport affinity across the cell and inner mitochondrial membrane. Future work is warranted to elucidate the physiological significance of these differences.

Our discovery that cellular BCKA uptake requires the plasma membrane monocarboxylate transporters MCT1 and MCT2 adds another critical regulatory layer to systemic homeostasis, one with immediate implications for ongoing drug development programs. MCT1 and MCT2 are currently major therapeutic targets in oncology, where small-molecule inhibitors (such as AR-C155858 and related clinical-stage compounds) are deployed to disrupt lactate and pyruvate fueling in highly glycolytic tumors (*25, 26*). Our findings reveal that these pharmacological interventions carry an unappreciated metabolic footprint: acute MCT1/2 inhibition by AR-C155858 in healthy mice rapidly compromises the systemic clearance of KIV and KIC, leading to a profound, selective accumulation of circulating BCKAs. This off-target effect must be carefully evaluated in clinical trial designs, as systemic BCKA elevations are linked to cardiometabolic disease (*27, 28*). However, strategically blocking cellular BCKA transport could hold independent therapeutic value for conditions characterized by pathologic BCKA hypercatabolic states or specific tumor dependencies. Because certain malignancies exploit BCKA-derived carbon networks to sustain lipid synthesis, the TCA cycle, DNA methylation, and to silence tumor laden immune cells, targeted MCT1/2 inhibition may represent a viable strategy to strip these cells of tumor promoting BCKA entry (*29–32*). Moreover, in contexts where a local reduction of BCKA oxidation is desired to maximize peripheral BCAA recycling—such as cachexia or protein-wasting states (*1*), the pharmacological modulation of cellular BCKA entry presents a clear alternative to direct enzymatic inhibition.

While our work provides the first identification of a substrate imported by SLC25A35 into mitochondria, Yamamuro *et al.* (*33*) recently reported a separate role for SLC25A35 in the mitochondrial export of phosphoenolpyruvate (PEP) to support adipose glyceroneogenesis. Viewed together, Yamamuro’s findings raise the possibility that counter exchange between PEP and BCKA via SLC25A35 might be a mechanism underlying part of the frequently reported crosstalk between the BCAA metabolic pathway and lipid metabolism (*34*). In line with this notion, Yamamuro’s untargeted mass spectrometry (MS)-based analysis of subcellular fractions from adipocytes lacking SLC25A35, detected accumulation of KIV in the cytosol in parallel to PEP accumulation in the mitochondria (*33*). Given that the number of mitochondrial metabolites far outweigh the 53 members of the SLC25 family of mitochondrial carriers (*18, 20*), it stands to reason that we are just scratching the surface of the transport function of SLC25A35 with BCKA and PEP. However, in conflict with our findings concerning BCKA, Yamamuro *et al.* presented data from LC-MS based proteoliposome assays using [^13^C]-labeled KIC and KIV in which they were unable to measure transport of BCKA by SLC25A35. Ultimately, they concluded that the effects of SLC25A35 deletion on adipocyte KIV were likely secondary metabolic effects of the primary action of SLC25A35 on PEP (*33*). This apparent disagreement with our findings, can be directly attributed to flawed analytical methodology used by Yamamuro to detect intra-liposomal BCKA. It is well documented that BCKA are highly labile and prone to rapid non-enzymatic degradation, thermal instability, and poor ionization in standard MS matrices unless structurally stabilized by chemical derivatization (*35, 36*). Crucially, Yamamuro’s un-derivatized workflow developed for PEP when applied to their proteoliposomes for assessment of BCKA transport failed to detect any of either the external ^13^C BCKA substrates or the 20 mM excess of unlabeled BCKA loaded into their liposomes (*33*). Thus, the appropriate conclusion to be drawn from these data is not inability of SLC25A35 to transport BCKA but rather that the analytical method used is invalid. By contrast, our use of radiolabeled substrate tracers provided an unambiguous, readout of direct transport activity. These methodological distinctions underscore the critical importance of ensuring appropriate substrate preservation when evaluating kinetic parameters for volatile metabolites.

Ultimately, our studies establishing SLC25A35 and the MCT axis as sequential gatekeepers of BCKA catabolism yield critical implications for human cardiometabolic diseases. Chronically elevated systemic BCAA and BCKA levels are robust, independent biomarkers for insulin resistance, type 2 diabetes, metabolic-dysfunction associated steatotic liver disease, and cardiovascular disorders (*27, 28*). Therapeutic interventions have historically targeted the enzymatic active state of BCKDH, primarily via small-molecule inhibitors like BT-2 to block the inhibitory kinase, BDK (*16*). However, our finding that BT-2 rapidly drives down systemic BCKAs—whereas hepatocyte-specific BDK deletion leaves blood BCKA pools entirely untouched—suggests that the therapeutic efficacy of these inhibitors relies on extrahepatic sites of action or BCKDH-independent effects such as BDK-mediated regulation of ATP-citrate lyase (*15*). Because SLC25A35 is naturally maintained at low, rate-limiting levels in the liver, simply upregulating hepatic BCKDH activity is ineffective for BCKA lowering. By shifting the dominant regulatory determinants of BCAA catabolism from the abundance of active dehydrogenase complex to inner mitochondrial membrane transport, our findings rewrite the fundamental rules governing systemic BCAA metabolic flux and provide new targets for BCKA-related therapeutics.

## ACKNOWLEDGEMENTS

Figures 1G, 2A, 2J, and 4F were created using BioRender.com. The authors thank Marlene J. Violette (Duke Proteomics and Metabolomics Shared Resource) for technical assistance.

## AUTHOR CONTRIBUTIONS

Conceptualization: YD, PJW, RWM, ZGH, IK

Methodology: YD, JMW, IK, MWF, GFZ, OI

Investigation: YD, JMW, IK, NM, SV, SF, EA, WH, MF, OI, GFZ

Visualization: YD, JMW, IK, NM, SV, WH, MF, GFZ

Funding acquisition: PJW, RWM, JMW, ZGH

Project administration: PJW, RWM, ZGH

Supervision: YD, PJW, RWM, ZGH

Writing – original draft: YD, PJW

Writing – review & editing: YD, RWM, PJW, IK, ZGH

## DATA, CODE, AND MATERIALS AVAILABILITY

All data are available in the main text and/or supplementary materials.

## STAR METHODS

### Key resources table

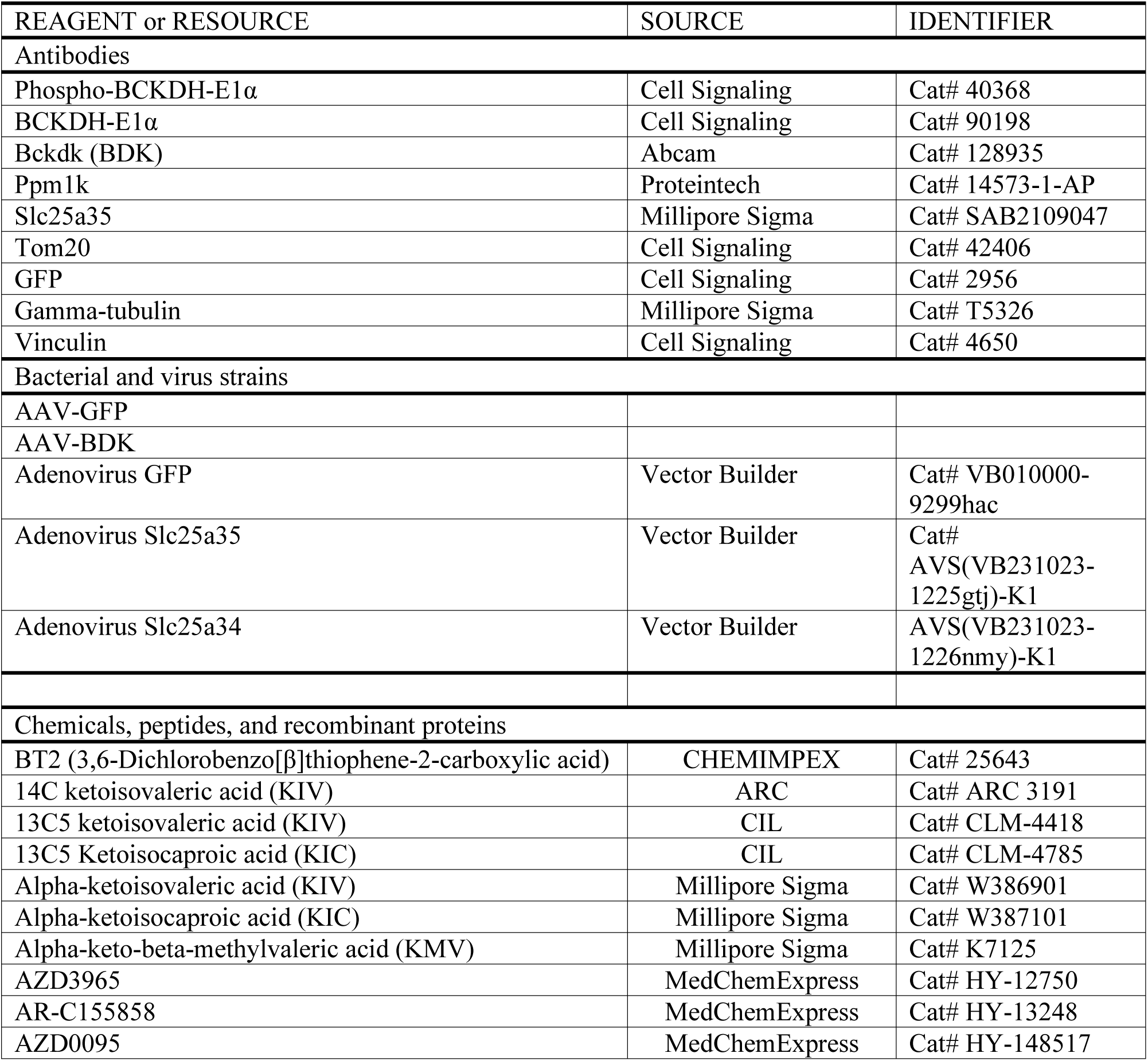
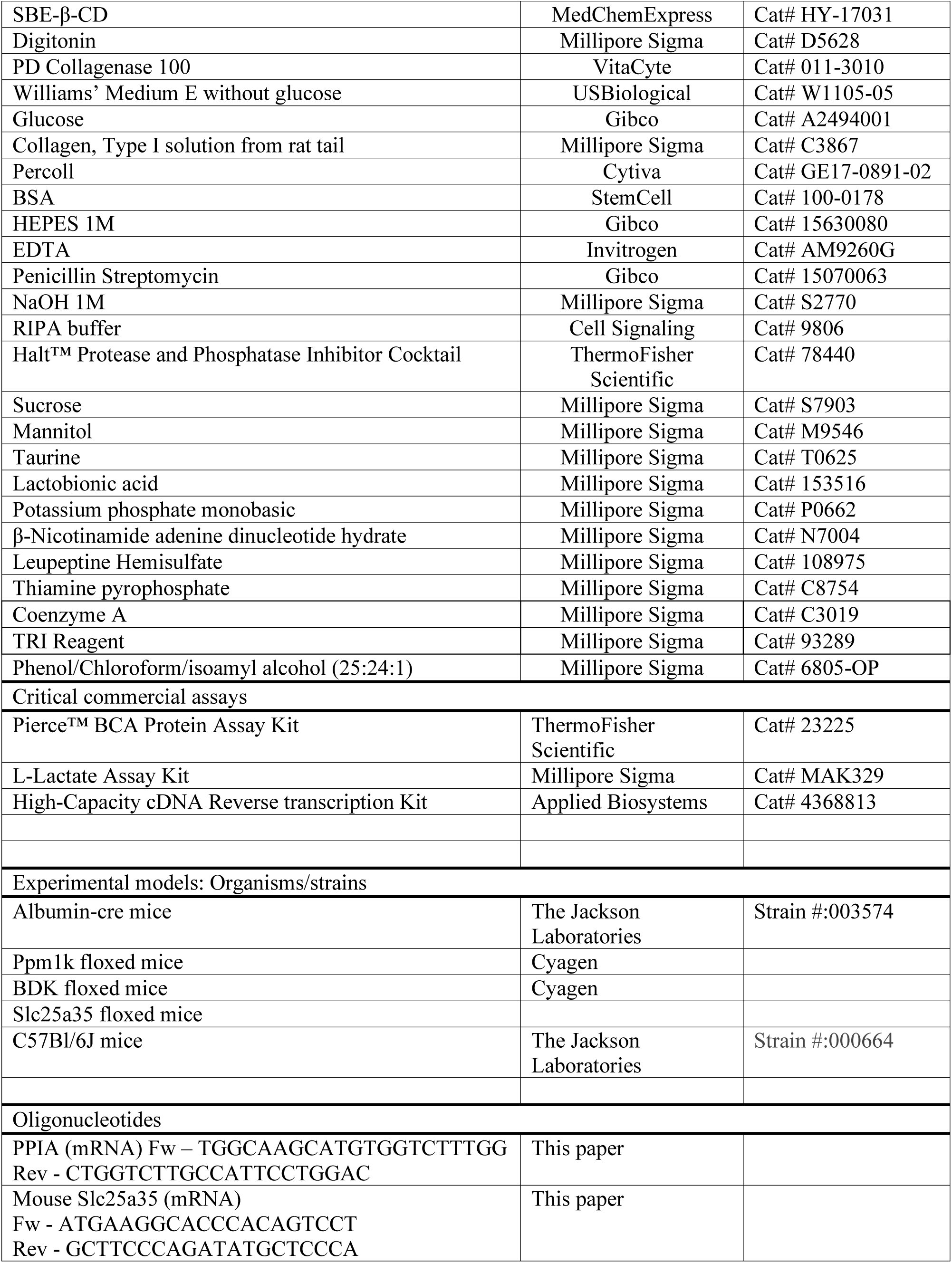
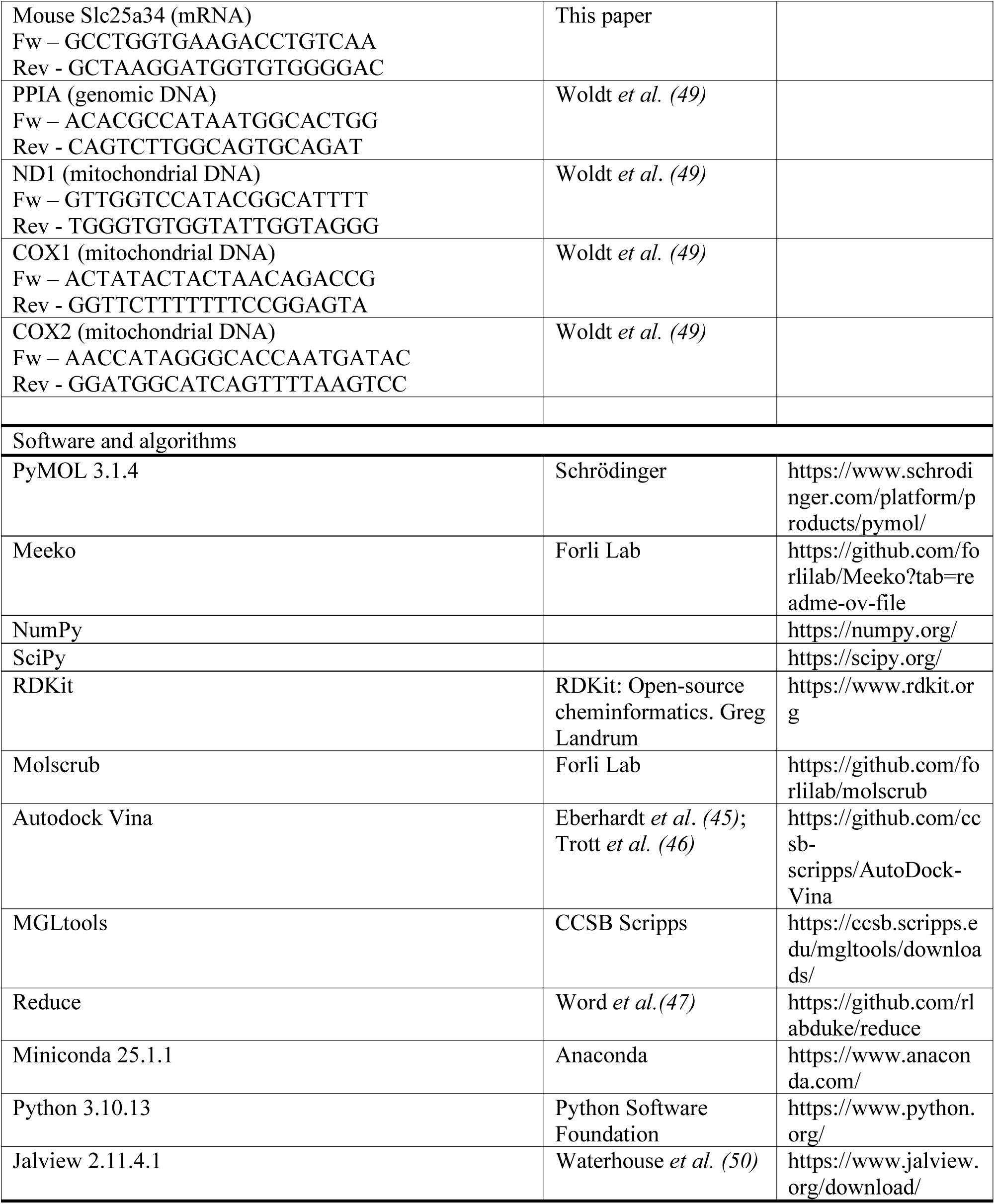

### Animals

All experiments involving mice were conducted in accordance with the guidelines established by the Duke Institutional Animal Care and Use Committee. Mice were maintained on a C57BL6 background and housed in a 12-hour dark-light cycle facility, with unrestricted access to water and food. Bckdk (BDK) and Ppm1k-floxed mice were generated by Cyagen and crossed with Albumin-Cre mice (The Jackson Laboratory, #003574), to generate liver specific KO animals.

Overexpression of BDK in the liver was achieved in 12-week-old male mice after a single retro-orbital injection with AAV8-CMV-BDK or AAV8-CMV-GFP (7×10^9^ viral particles per animal). Mice were euthanized 8 weeks post infection and expression of BDK in the liver was confirmed by WB. Adenoviral overexpression of SLC25A35or GFP control was achieved in 12-week-old male mice by a single 50µl retro-orbital injection of Ad-CMV-SLC25A35 or Ad-CMV-GFP adenovirus at a dose of 1×10^9^ IFU per animal.

Studies with SLC25A35 mice were performed with approved protocols from The Danish Animal Experiments Inspectorate (permit number: 2023-15-0201-01386) and the University of Copenhagen (project number: P23-336). Mice with constitutive whole body *Slc25a35* knockout 6N(Cg)-Slc25a35^tm1.1(KOMP)Vlcg^/J (SLC25A35 KO) were purchased from Jackson Laboratories (Stock No: 13834A-A6) in a heterozygous state. Littermate wild-type mice were used as controls in the experiments. Mice were maintained on a C57BL6 background and housed in a 12-hour dark-light cycle facility, with unrestricted access to water and food. Male mice acclimated to room temperature were singly housed for two weeks and dissected at 14-18 weeks of age. Blood plasma samples were collected by cardiac puncture under isoflurane anesthesia, and mice were sacrificed afterward.

### BCKA tolerance tests

For MCT inhibitor studies, 12-week-old male mice were fasted for 4 hours and intraperitoneally injected with 5 mg/kg AR-C155858 (a MCT1/2 inhibitor) in 10% DMSO, 20% SBE-β-CD, in saline. 2 hours after receiving the inhibitor, mice were intraperitoneally injected with a combination of all three BCKAs at 100 mg/kg. Blood samples were collected at the tail tip before and 2 hours after AR-C155858 injection, as well as at 5, 15, and 45 minutes after BCKA injection. The blood was then centrifuged for 15 minutes at 5,000 x g, and the plasma was snap frozen. For Ad-SLC25A35 studies, the BCKA tolerance test was conducted 5 days following the administration of the adenovirus following a 5 hour fast as described above.

### Mouse primary hepatocytes isolation, culture and treatments

Primary hepatocytes were isolated from 12 to 20-week–old male mice using the two-step collagenase perfusion method, as previously described (*37*). Briefly, the liver was perfused through the inferior vena cava to wash out blood and chelate calcium, followed by perfusion with a buffer containing 0.6 g/mL collagenase (PD collagenase 100, 011-3010, Vitacyte) and kept on ice in 10 mL of William’s E media supplemented with 11 mM glucose, 0.1% BSA, 100 nM dexamethasone and 1 % of penicillin/streptomycin (maintenance media), until dissociation. After dissociation, hepatocytes were washed in maintenance media, centrifuged for 2 minutes at 50 x g and then passed through a 70 μm filter. Enrichment of live viable primary hepatocytes was then performed using 45% Percoll in maintenance media and centrifuged for 10 minutes at 400 x g. The pellet was resuspended and hepatocytes were counted and seeded at a density of 450,000 cells per well of a 6-well plate, coated with collagen. After 2 to 3 hours, media was refreshed. All experiments were conducted the following day.

For SLC overexpression studies in primary hepatocytes, 2 hours post isolation, primary hepatocytes were infected with either Ad-CMV-SLC25A34 or Ad-CMV-SLC25A35 at multiplicity of infection (MOI) of 25. The next day, approximately 16 hours after transfection, primary hepatocytes were used to assess the oxidation of 1-[^14^C]-KIV.

### Mitochondria isolation

After euthanasia, a piece of mouse liver was kept on ice in MIR05 buffer (20 mM HEPES, 10 mM KH2PO4, 110 mM sucrose, 20 mM taurine, 60 mM K-lactobionate, 0.5 mM EGTA, 3 mM MgCl2·6H2O, 1 g/L of BSA fatty acid free). 200 mg of liver were weighed, minced and transferred into a glass Potter-Elvehjem. Homogenization was achieved by 10 strokes with a PTFE pestle in 2 mL of mitochondria isolation buffer (225 mM Mannitol, 75 mM Sucrose, 0.2 mM EDTA). The homogenates were then transferred to a 2 mL microcentrifuge tube and centrifuged at 600 x g for 10 minutes at 4°C. The pellet was discarded and the supernatant was centrifuged a second time for 10 minutes at 6,200 x g for 10min at 4°C. The mitochondria pellet was resuspended with 500 μL MIR05 buffer.

### 1-[14C]-KIV oxidation assays

For assays in primary hepatocytes, the day following their isolation, mouse primary hepatocytes were incubated in 1.25 mL of culture media and 50 μL of media containing 20 mM HEPES, 1 μL of 1-[^14^C]-KIV and 100nmol of cold KIV were added per well. Plates were sealed with 2 layers of parafilm and incubated at 37°C and 5% CO_2_ for 45 minutes. After incubation, 1mL of culture media was transferred into a polystyrene test tube, on ice, along with a CO_2_ trap (200 μL NaOH 1 M) and the tube was tightly capped. The media was then acidified by injection of 100 μL of 70% perchloric acid and tubes were left on an orbital shaker for 1 hour. ^14^CO_2_ contained in the trap was counted on a Beckman Coulter LS6500 liquid scintillation counter.

For isolated mitochondria, the 1-[^14^C]-KIV oxidation assay was performed with 50 μL of freshly isolated liver mitochondria, in duplicate, 300 μL of MIR05 containing 0.1 μCi of 1-[^14^C]-KIV and 100 nmol of cold KIV, as well as the CO_2_ trap (200uL NaOH 1 M) in a polystyrene test tube. All tubes were then capped and incubated for 30 minutes at 37°C in a shaking water bath. After incubation, 100 μL of 70% perchloric acid were added to the reaction and tubes were incubated on an orbital shaker for 1 hour. ^14^CO_2_ contained in the trap was counted on a Beckman Coulter LS6500 liquid scintillation counter.

For assays in liver lysates, 1-[^14^C]-KIV oxidation was performed with 25 mg of frozen mouse liver, pulverized in liquid nitrogen and homogenized using a QIAGEN TissueLyser II in 250 μL of ice-cold buffer H (30 mM KPi pH 7.5, 3 mM EDTA, 5 mM DTT, 1mM KIV, 3% FBS, 5% Triston-X-100, 1uM Leupeptin) or 450,000 mouse primary hepatocytes lysed in 125 μL buffer H. After centrifugation at 10,000 xg for 10min, 50 μL of liver homogenates were added to 300 μL of buffer A (50 mM HEPES pH 7.5, 30 mM KPi pH 7.5, 0.4 M CoA, 3 mM NAD+, 5% FBS, 2 mM Thiamine pyrophosphate, 2 mM MgCl2, 0.1 μCi 1-[^14^C]-KIV), in a polystyrene tube containing a CO_2_ trap. After incubation at 37°C in a shaking water bath (30 minutes for tissue homogenates, 1 hour for lysed primary hepatocytes), the reaction is acidified by injection of 70% perchloric acid and tubes are incubated on an orbital shaker for 1 hour. ^14^CO_2_ contained in the trap was counted on a Beckman Coulter LS6500 liquid scintillation counter.

### Immunoblotting

Lysates were prepared in RIPA lysis buffer (Cell Signaling Technologies) containing a protease and phosphatase inhibitor cocktail (Thermo Fisher Scientific). Protein concentration was assessed using Pierce BCA assay kit (Thermo Fisher Scientific) and 20 to 50 μg of protein were loaded onto 4-15% Tris-Glycine gels (Bio-Rad), subjected to SDS-PAGE and transferred on PVDF membranes. Blocking, primary and secondary antibody incubations were all performed in “Fish buffer” (0.45% fish gelatin, 0.1% N-Z-amine A and 0.02% sodium azide, in TBS). Primary antibodies were used at 1:2000 and secondary antibodies at 1:10000. All antibodies used are listed in the Key Resources Table. Membranes were scanned using a Li-Cor Odissey CLx.

### LC-MS/MS Analysis of Branched-Chain Keto Acids (BCKA), Branched-Chain Amino Acids (BCAA), and Hydroxybutyrates (HB)

10 µl of plasma or 200 µl of tissue homogenate prepared in 3M perchloric acid (PCA) at 50 mg of wet tissue/ml were spiked with isotopically labeled internal standards KIC-d3, KIV-13C5 (Cambridge Isotope Laboratories, MA, USA), KMV-d8 (Toronto Research Chemicals, Canada), leucine-d3, isoleucine-13C6, valine-d8, 2-HIB-d6, 2-HB-d3, and 3-HB-d4 (CDN Isotopes, Canada). Plasma was precipitated with 150 µl of 3M PCA, and both tissue and plasma samples were centrifuged for 5 min. at 14,000g. 200 µl of 25 M o-phenylenediamine (OPD) in 3M HCl was added to the supernatants and the samples were incubated at 80°C for 20 minutes. Metabolites were extracted with ethyl acetate as previously described (*38–40*). The extracts were dried under nitrogen, reconstituted in water, and analyzed on a Waters Xevo TQ-S triple quadrupole mass spectrometer coupled to a Waters Acquity UPLC system (Milford, MA). The analytical column (Waters Acquity UPLC HSS T3, 1.8 μm, 2.1 × 100 mm) was maintained at 35°C. 2 μl of the sample was injected onto the column and eluted at a flow rate of 0.4 ml/min. 0.1% formic acid in water was used as a mobile phase A and acetonitrile was used as a mobile phase B. MRM transitions for BCKA and BCAA were monitored in the positive ion mode and for hydroxybutyric and hydroxyisobutyric acids in the negative ion mode. Metabolite concentrations were computed using external calibration constructed from a serial dilution of the standards (Sigma, MO, USA) in dialyzed fetal bovine serum.

### Stable isotope-resolved metabolite profiling

Tissues were harvested 10 min after the U-[^13^C]-KIV injection and immediately freeze-clamped in liquid nitrogen and stored at −80 °C until further analysis. Frozen tissues were pulverized under liquid nitrogen using a mortar and pestle. Metabolites were then extracted using sequential 500 μl additions of −20 °C MeOH, chilled water, and chloroform. After each addition, tissue lysates were prepared with a Tissue Lyser (Qiagen) for 60 s at 30 Hz. Tissue extracts were then centrifuged at 4 °C and 14,400 × *g* for 20 min and the clarified aqueous phase was transferred to a fresh Eppendorf and stored in −80 °C until processing by GC-MS analysis.

For GC-MS analysis, the extracted tissue metabolites were dried under N_2_ gas-flow at 37 °C using an evaporator and organic acids were derivatized via methoximation and silylation as previously described (*9*). Briefly, metabolites were resuspended in 25 μl of methoxylamine hydrochloride (2% (w/v) in pyridine) and incubated at 40 °C for 90 min on a heating block. After brief centrifugation, 35 μl of MTBSTFA + 1% TBDMS was added and the samples were incubated at 60 °C for 30 min. GC-MS analysis was performed on an Agilent 7890B GC system equipped with an HP-5MS capillary column connected to an Agilent 5977 A Mass Spectrometer. Mass isotopomer distributions were obtained by integration of ion chromatograms and corrected for natural isotope abundances. Metabolites monitored were 3-HIB (*m/z*: 275–280) and citrate (*m/z* 459–467).

### RNA extraction, reverse transcription and real time quantitative PCR analysis

Total RNA was isolated from tissue or cells using TRI Reagent (Millipore Sigma). cDNA was generated from 1 µg of RNA, using the high-capacity reverse transcription kit (Applied Biosystem, Life technologies). Real time QPCR was performed on a Quantstudio 6 Pro (ThermoFisher Scientific), using specific primers (see Resource table). Analysis was performed with the ddCt method.

### SLC25A35 co-expression analysis

The GEO database was manually searched for Affymetrix Mouse Genome 430 microarray datasets in which *Slc25a35* was detected and significantly up or downregulated under at least one experimental condition. Four datasets representing distinct tissues were identified: (i) GDS3533 (*41*), virgin mammary gland; (ii) GDS4883 (*42*), liver; (iii) GDS4856 (*43*), skeletal muscle (soleus); and (iv) GDS3768 (*44*), gonadal white adipose tissue. For each dataset, genes whose expression was regulated in the same direction as *Slc25a35* were extracted using a significance threshold of p < 0.01, without applying a fold-change cutoff. Genes common to all four datasets were then identified, resulting in a set of 56 genes. Pathway enrichment analysis using KEGG pathway annotations was performed on this list of 56 genes.

### Molecular docking

The hydrated and flexible docking simulations were performed in Python using Autodock Vina (*45,46*). AlphaFold PDB file containing the receptor coordinates for Slc25a35 (AF-Q3KQZ1-F1-v4) were first modified with the reduce script (*47*) ($ reduce -BUILD receptor.pdb), adding all missing hydrogen atoms. PDBQT file and grid parameters of the receptor were then generated using the python package Meeko ($ mk_prepare_receptor.py) with the following grid coordinates (x y z; box size), 1.79 2.256 −0.083; 40 40 40 (*48*). K24, K31, K127, R175 and R276 were specified to be flexible residues. Affinity maps and water map were finally computed with autogrid4 and the python Vina package mapwater.py, respectively. Ligands coordinates were obtained through PubChem (sdf files, 3D structures) and hydrogen atoms were first added with the python package Molscrub ($ scrub.py -I ligand.sdf −0 ligandH.sdf). The meeko script ($ mk_prepare_ligand.py -i ligandH.sdf -o ligand.pdbqt -w) was then used to add explicit water molecules to the ligands and generate the PDBQT files. The Docking affinities were calculated using Autodock Vina with Autodock 4 forcefield and the exhaustiveness set at 32, following Vina hydrated docking protocol (https://autodock-vina.readthedocs.io/). The computed docking was visualized in the software PyMOL (Schrodringer), showing residues implicated in the matrix salt bridges of the transporter, as well as interacting with the ligands.

### Proteoliposomes preparation and transport assay

Human SLC25A35 protein was synthesized by Synthelis using an E.Coli cell free protein synthesis system, in the presence of liposomes. Proteoliposomes were isolated on a sucrose gradient, pelleted at 10,000 x g and resuspended in 20mM HEPES buffer. Transport was assayed by incubating the proteoliposomes or control empty liposomes in HEPES buffer containing 2.5 mM cold KIV and 0.1 uCi 1-14C-KIV, for 45 minutes at 37°C. Transport in the proteoliposomes was then stopped by the addition of ice-cold HEPES wash buffer. Following washing with ice-cold HEPES buffer, proteoliposomes were pelleted by gentle centrifugation for 10 minutes at 10,000 x g. The pellet was then resuspended in a mixture of chloroform: methanol (2: 1), to liberate the imported substrate. Samples were then transferred into scintillation liquid and counted on a Beckman Coulter LS6500 liquid scintillation counter.

### Quantitative proteomic analysis

15 µgs of mitochondrial lysates or cytosol (n=3 per group) were adjusted to 5% SDS and 50 mM triethylammonium bicarbonate, pH 8.5 (TEAB) followed by denaturation and reduction with 10 mM DTT at 80 °C for 10 min, alkylation with 25 mM iodoactemide at room temp for 30 min and digestion with 1.5 µg of sequencing grade modified trypsin (Promega) for 2 h at 47 °C using an S-trap micro device (Protifi). Eluted peptides were lyophilized and reconstituted in 30 μL of 0.1% formic acid. A study pool QC (SPQC) was created by combining equal volumes of all samples, and 2 μL of each sample and the SPQC was loaded on Evotips.

LC/MS/MS was performed on each sample and three replicates of an SPQC pool, using an Evosep One LC and a Thermo Orbitrap Astral. Briefly, the Evosep used a 30 sample-per-day method with a Pepsep 15 cm x 150 µm column. The MS analysis used a 240,000 resolution precursor ion (MS1) scan and data-independent MS/MS with 4 m/z isolation windows from 380-980, a maximum ion transfer time of 6 ms and normalized collision energy of 20 ms. Data was processed in Spectronaut 19 (19.0.240606.62635; Biognosys). Direct-DIA searches used a *Mus musculus* database (downloaded from Uniprot on 4/08/2022), trypsin/P specificity with up to 2 missed cleavages, fixed carbamidomethyl (Cys) modification and variable oxidation (Met) and protein N-terminal acetylation. Data was annotated at a 1% precursor and protein group false discovery rate (q-value). Localization normalization and MaxLFQ protein abundance calculations were performed in Spectronaut. Raw and processed data have been uploaded to the ProteomeXchange consortium (PXD078778) via the MassIVE repository (MSV000101947) and can be accessed via the web interface with reviewer login “MSV000101947_reviewer” and password “BCKA” or via ftp://MSV000101947@massive-ftp.ucsd.edu with username “MSV000101947” and password “BCKA”.

## Supplementary Materials

Figs. S1 to S4

