## Supplementary Materials for "Branched-chain α-keto acid metabolism is controlled by SLC25A35-mediated mitochondrial import"

**The PDF file includes:**

Figs. S1 to S4

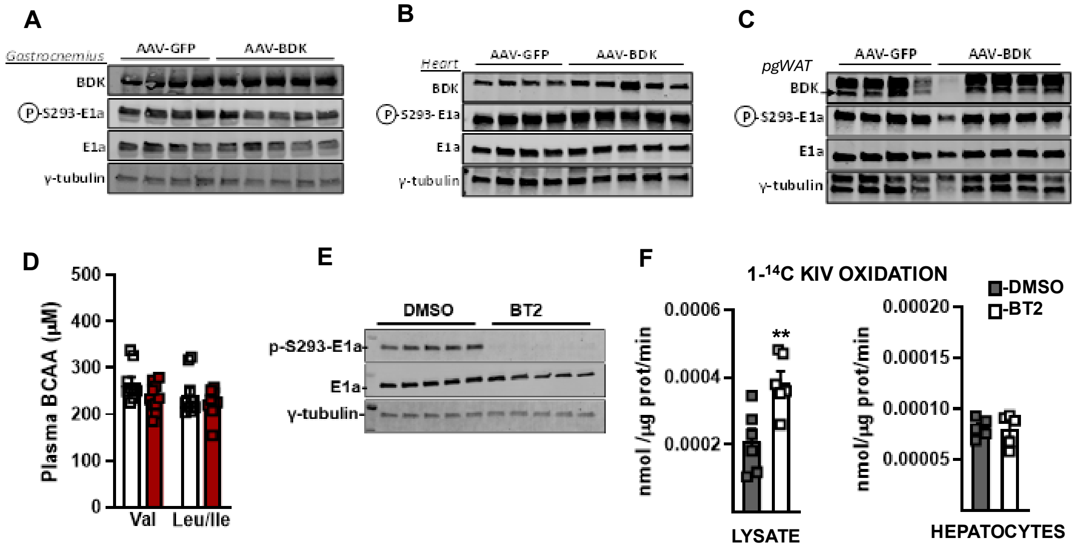


**Fig. S1.**

Western Blot analysis of BDK, phosphorylated and total E1a and γ-tubulin in (**A**) gastrocnemius, (**B**) heart, and (**C**) epididymal adipose tissue from mice injected with AAV-CMV-BDK or GFP. (**D**) Circulating BCAA in BDK overexpressing mice and controls. **(E)** Western Blot analysis of phosphorylated and total E1a as well as (**F**) 1-[$^{14}$C] KIV oxidation rate in lysate or in live mouse primary hepatocytes treated with 50µM BT2 for 2h. Statistics: (F) Unpaired two-tailed t-test, ** p <0.01.

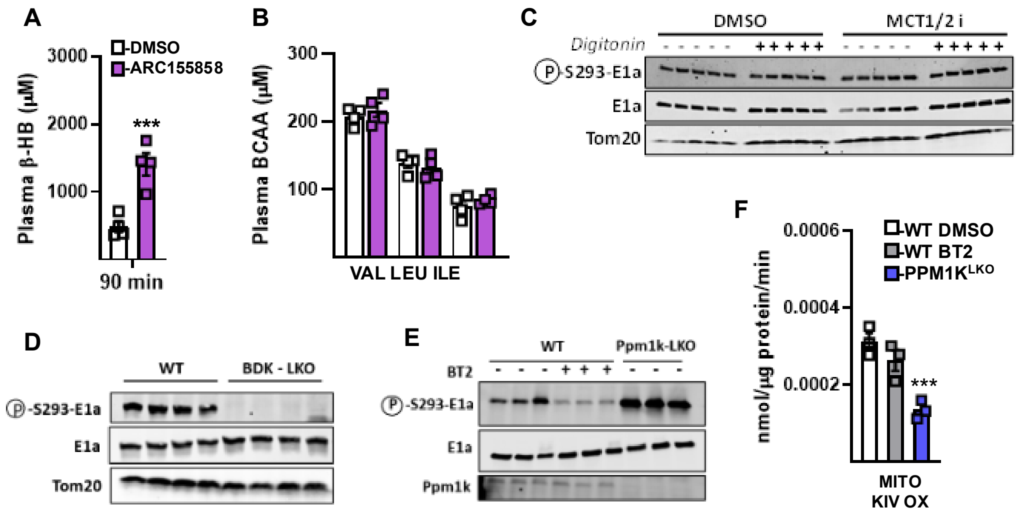


**Fig. S2.**

(**A**) Circulating β-HB and (**B**) circulating BCAA in mice treated with MCT1/2 inhibitor (ARC-155858; 5mg/kg) for 90min. (**C**) Western Blot analysis of phosphorylated E1a, total E1a and Tom20 in primary hepatocytes treated with digitonin (7.5ug/mL) and MCT1/2 inhibitor (AR-C155858; 250nM). (**D**) Western Blot analysis of phosphorylated E1a, total E1a and Tom20 in isolated liver mitochondria from BDK LKO mice or control littermates. Ppm1k fl/fl mice were treated with $20mg.kg^{-1}$ BT2 or DMSO for 5 consecutive days and compared to littermate $PPM1K^{LKO}$ mice for (**E**) hepatic levels of phosphorylated E1a, total E1a and Ppm1k and (**F**) rate of oxidation of 1-[$^{14}$C] KIV in isolated liver mitochondria. Statistics: (A) Unpaired two-way t-test, *** p <0.001. (F) One-way ANOVA with Dunnett's multiple comparisons test, ** p<0.01.

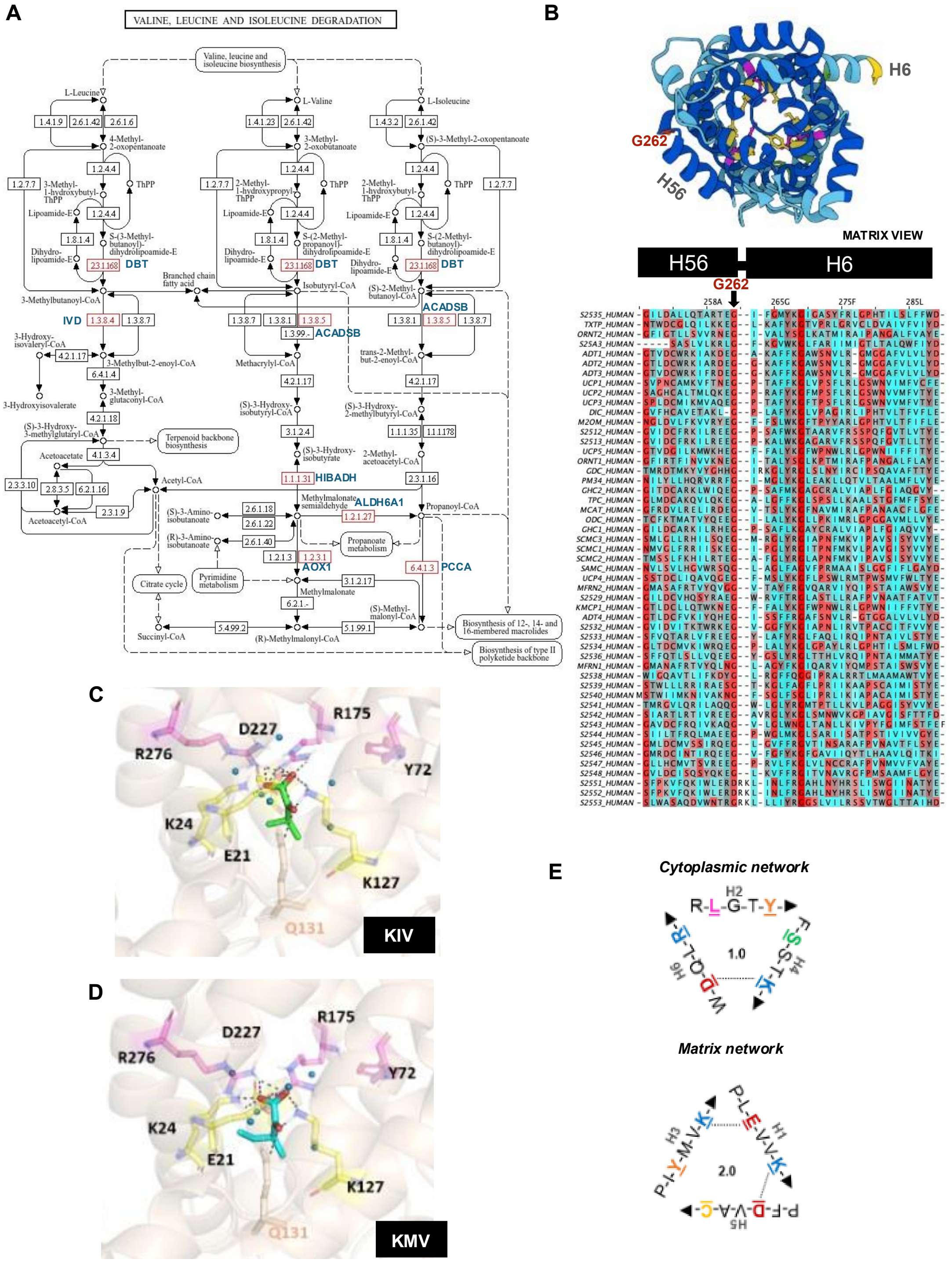


**Fig. S3.**

**(A)** KEGG map of the valine, leucine, and isoleucine degradation pathway with SLC25A35 coregulated genes shown in red alongside gene names in blue. **(B)** Top panel, cartoon representation of human SLC25A35, viewed from the matrix side of the mitochondrial membrane, with glycine residue G262 highlighted in red. Bottom panel, multiple protein sequence alignment generated in Jalview, using Clustalw, of 51 members of human SLC25A carriers, centered around residue G262. Color scheme showing turn propensity. (**C-D**) Lateral view (c-state) of the simulated hydrated-flexible docking of human SLC25A35 with KIV and KMV respectively, metabolites are represented as sticks. Polar interactions are represented with dashed grey lines. Other non-flexible residues found to form interactions with KIV and KMV are shown in wheat. (**E**) Relative cytoplasmic (left) and matrix (right) salt bridge network strength of SLC25A35.

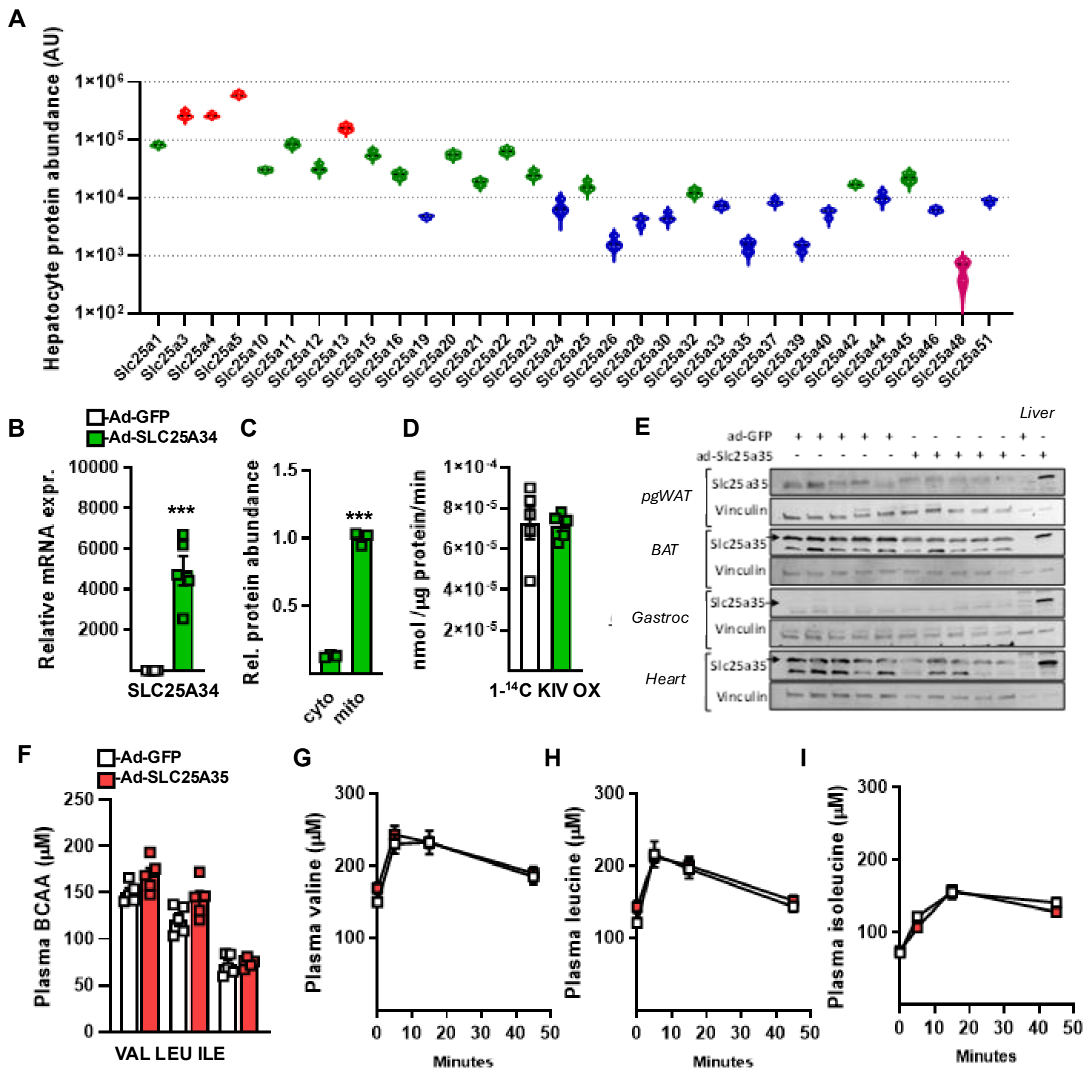


**Fig S4.**

(**A**) Protein abundance of SLC25A members measured by mass spectrometry in primary hepatocytes (n=3). (**B**) Relative mRNA expression of SLC25A34 in primary hepatocytes transfected with Ad-CMV-SLC25A34 or Ad-CMV-GFP (n=5 per condition). (**C**) Subcellular localization of SLC25A34, measured by mass spectrometry, in cytosolic and mitochondrial fractions of primary hepatocytes transfected with Ad-CMV-SLC25A34 (n=3). (**D**) Rate of 1-[$^{14}$C]-KIV oxidation in primary hepatocytes transfected with Ad-CMV-SLC25A34 or Ad-CMV-GFP (n=5 per condition). (**E**) Western Blot analysis of SLC25A35 protein level in perigonadal white adipose tissue (pgWAT), brown adipose tissue (BAT), gastrocnemius and heart in comparison to the liver of mice receiving a single injection of Ad-CMV-SLC25A35 (n=5 per group). (**F**) Circulating BCAA in mice overexpressing SLC25A35 or GFP in the liver. (**G**) Valine, (**H**) Leucine, and (**I**) Isoleucine concentration in plasma during the BCKA tolerance test. *Statistics*: All values are expressed as means ± SEM. (B, C) Unpaired two-tailed t-test, *** $p<0.001$.
